# Comparative genomic assessment of the *Cupriavidus necator* species for one-carbon based biomanufacturing

**DOI:** 10.1101/2025.03.26.645245

**Authors:** Magnus G. Jespersen, Emil Funk Vangsgaard, Mariana Arango Saavedra, Stefano Donati, Lars K. Nielsen

## Abstract

The transition from a petroleum-based manufacturing to biomanufacturing is an important step towards a sustainable bio-economy. In particular biotechnological processes which use one carbon (C1) compounds as feedstock represent an interesting avenue. Many bacterial species evolved naturally to thrive on such compounds, among them *Cupriavidus necator,* which has been studied in the past due to its range of metabolic capabilities in utilization and production of compounds of interest. *Cupriavidus necator* strain H16 is the reference laboratory strain for this species and by far the most extensively studied. In contrast, research efforts and genomic characterization of other strains within this species have been limited and sporadic. Therefore, the genomic diversity and full metabolic potential across the broader species remain poorly understood.

In this work, we collected publicly available genomes along with newly sequenced ones. From a collection of 44 genomes we curated a final collection of 22 genomes deemed to be *C. necator*. We examined hallmark metabolic functions, including carbon dioxide fixation, formate assimilation, and hydrogen utilization. We identified methylation motifs and restriction modification systems. Finally, strains ATCC 25207, TA06, and 1978 are proposed as candidate strains of interest based on their genomic make-up and observations from literature. This work provides a comprehensive genomic resource for the *C. necator* species, facilitating its development as a biomanufacturing platform and advancing our understanding of its metabolic diversity and potential applications.

**Importance:** The green transition from petroleum-based manufacturing to a biomanufacturing economy requires new solutions. One of these is the development of bacterial platforms for production of valuable chemicals, in particular from CO2 and derived molecules. In this context, *Cupriavidus necator* is one of the most studied platform strains, although the genomic potential of this species has been seldom explored. This work provides a curated set of *C. necator* genomes, collected from public repositories and additional newly sequenced genomes. Through multiple analyses, *C. necator* ATCC 25207, TA06, and 1978 are proposed as strains of interest for further use. These findings lay the groundwork for advancing *C. necator* as a key organism in sustainable biomanufacturing, highlighting its potential for efficient carbon fixation, formate assimilation, and hydrogen utilization. The identification of methylation motifs and restriction modification systems further supports genetic engineering efforts, making these strains highly valuable for future biotechnological applications.

## Introduction

Biomanufacturing is seen as an alternative to the current petroleum-based manufacturing and is a growing ambition of both private and public actors (1). One path for biomanufacturing involves the fixation of one carbon (C1) based compounds, such as methanol, formaldehyde, formate, and carbon dioxide. C1 based biomanufacturing is viewed as a way of achieving a truly circular manufacturing loop, as it upgrades waste and greenhouse gas to value-added goods (2). A key component in a C1 based biomanufacturing framework are microbes capable of utilizing C1 compounds. Two major strategies to obtain such microbes are being explored. One strategy takes current model organisms like *Escherichia coli* and adapts these to de novo assimilate C1 compounds (3). The second strategy takes existing C1 assimilating microorganisms and optimize these for manufacturing.

*Cupriavidus necator* has received attention through the years due to the ability of some strains to grow autotrophically on CO_2_ and molecular hydrogen. Initially known under the umbrella of the genus of *Hydrogenomonas* (4), along with other species names: *Hydrogenomonas eutrophus, Alcaligenes eutrophus, Ralstonia eutropha,* and *Wautersia eutropha.* The wide range of carbon sources utilized by *C. necator* combined with a versatile metabolism able to produce Polyhydroxyalkanoates (PHAs) have kept this organism of high interest for biomanufacturing applications (5). *C. necator* strain H16 was isolated in the 1960’s (4) and has established itself as the laboratory reference strain. H16 can assimilate molecular hydrogen as an energy source, allowing it to produce biomass from carbon dioxide and molecular hydrogen alone. Finally, *C. necator* H16 can also grow on formate, by oxidizing it into carbon dioxide to be fixed with the reducing power obtained from the oxidation.

As *C. necator* was first named *Hydrogenomonas eutropha* in the 1950’s (4), along with other hydrogen assimilating species, it has changed species designation multiple times. These species reclassifications have resulted in a scattered and confusing mesh of information about *C. necator*. In the genomic era, multiple *C. necator* genomes have been sequenced, with attempts to define genomic hallmarks for *C. necator* (5, 6). One proposed hallmark is to lower the usual threshold of 95% Average Nucleotide Identity for *same* species genomes to 90% for *C. necator*. Besides species identification, *C. necator* genomics has been used to yield insights into methylation patterns leveraged to increase transformation efficiency (7–9). Although multiple genomes of *C. necator* have been sequenced, no single study has curated and compared these genomes further.

In this work we expand the number of sequenced and complete *C. necator* genomes. In addition, we collect publicly available genomes, to produce a collection of curated *C. necator* genomes, excluding genomes with incorrect annotation. The collection is examined for metabolic traits of interest in *C. necator*. Finally, we identify methylation motifs for restriction modification systems in select strains, to enable higher transformation efficiency for future genetic engineering campaigns.

## Results

### Expanding the number of complete genomes of *Cupriavidus necator*

To expand the knowledge of *Cupriavidus necator* genomics, a collection of wild type isolates annotated as *Cupriavidus necator* was ordered from the Deutsche Sammlung von Mikroorganismen und Zellkulturen (DSMZ)(10–12). Genomes were sequenced using both short- and long-read technologies and assembled using hybracter (13). Eleven genomes were sequenced, nine of these were newly sequenced, with two, H850 (11) and H16 (12), being re-sequenced. All genomes were assembled to completion with closed chromosomes, chromids, and plasmids (Supplementary Table 1). Overall, we provide two improved and nine newly sequenced complete genomes of *Cupriavidus necator* strains as annotated by DSMZ.

### Assessment of *Cupriavidus necator* genomes

To supplement newly and improved genome sequences, we collected publicly available genomes that were either annotated as *C. necator* or *Cupriavidus* species (sp.), had been associated with *C. necator* in previous studies (5, 6), or annotated as *C. necator* in large assembly and species annotation efforts (14). This effort resulted in 30 prospective *C. necator* genomes from public sources (6, 15–28) (Supplementary table 1), and inclusion of three genomes of other species as outgroups (29–31). Outgroup isolates were selected based on a historical association with *C. necator*, *Cupriavidus tuberpinensis* strain JMP134, or close genomic association with *C. necator*, *Cupriavidus taiwanensis* strain LMG19424 and *Cupriavidus lacunae* strain S23.

A collection of 44 genomes, including those sequenced in-house, publicly available, and from known outgroups was assembled to be evaluated. Initial refinement of the genome collection was based on k-mer based taxonomic sequence classification (32) and evaluation of two average nucleotide identity analyses (ANI)(33, 34) (Supplementary Table 1). K-mer based taxonomic sequence classification highlighted 18 strains from public repositories as unlikely *C. necator* strains (*Cupriavidus necator* match < 2% of sequences). Of the 18 strains, 10 had very low ANI coverage (<40% to both *C. necator* H16 and *C. necator* N-1). This included the *C. pinatubunensis* outgroup representative, JMP134, which was included in further analyses.

Of the 8 strains, with >40% ANI coverage, 6 were associated with *C. taiwanensis*, including the outgroup representative LMG19424. Isolate KK10, was by k-mer analysis associated 100% to itself as a *Cupriavidus sp.*. Finally, the in-house sequenced Tfa17 was by k-mer analysis associated with *Cupriavidus oxalaticus*, with acceptable ANI coverage, ∼49% against H16 and ∼56% against N-1. Based on these observations, we keep LMG19424 as an outgroup representative of *C. taiwanensis* and KK10 and Tfa17 for further analyses. In summary, using a k-mer based species classification cutoff of 2%, ANI coverage cutoff against *C. necator* H16 and N-1 of 40%, and collapsing species into a single outgroup representative, we refined the initial genome collection from 44 genomes to 30 for further analyses (Supplementary Figure 1). Of these 30 strains, 9 genomes were the ones sequenced for the first time in this study (Supplementary Figure 1).

### Refining *Cupriavidus necator* strains using average nucleotide identity as a marker of species

The refined collection of 30 genomes were assessed in depth in terms of ANI coverage and identity percentage. Previously an ANI percentage cutoff of 90% has been proposed for *C. necator* to identify strains of the species (6). This is in contrast to the usual ∼95% ANI percentage threshold used for most species and derived from DNA-DNA hybridization (35). In a phylogeny of the 30 genomes, strains NH9 and *C. lacunae* S23 are placed on a monophyletic clade, that is the closest to the monophyletic *C. necator* clade (Figure 1A). The relationship between NH9 and S23 is further exemplified by their ANI (∼97%) and coverage (∼69%)(Figure 1B), indicating that the more regularly used ANI cutoff of 95% can be applied. This would classify strain NH9 as a *C. lacunae*. Similar to NH9, A5-1 seems to be misclassified as *C. necator* in public repositories. A5-1 and DTP0602 are around the ANI species cutoff (∼94%), indicating these isolates to be of the same species. The grouping of these isolates has previously been observed (6), confirming the annotation of A5-1 as a *C. necator* to most likely be a misannotation of species. Stains SHC2-3 and Tfa17 both classified as *C. necator* in public repositories and the DSMZ strain bank, respectively, both seem to be a species other than *C. necator*, as these are nestled among known outgroups in the ANI tree. The analysis of ANI Identity percentage leaves 22 genomes classified as *C. necator* with an ANI ≥94% and a coverage of ≥50% (Supplementary Figure 1). Based on ANI of the outgroups included in this analysis, *C. lacunae* seems to be a close relative of *C. necator*, and a key outgroup to include in future genomic analyses (Supplementary Figure 2).

**Figure 1.**
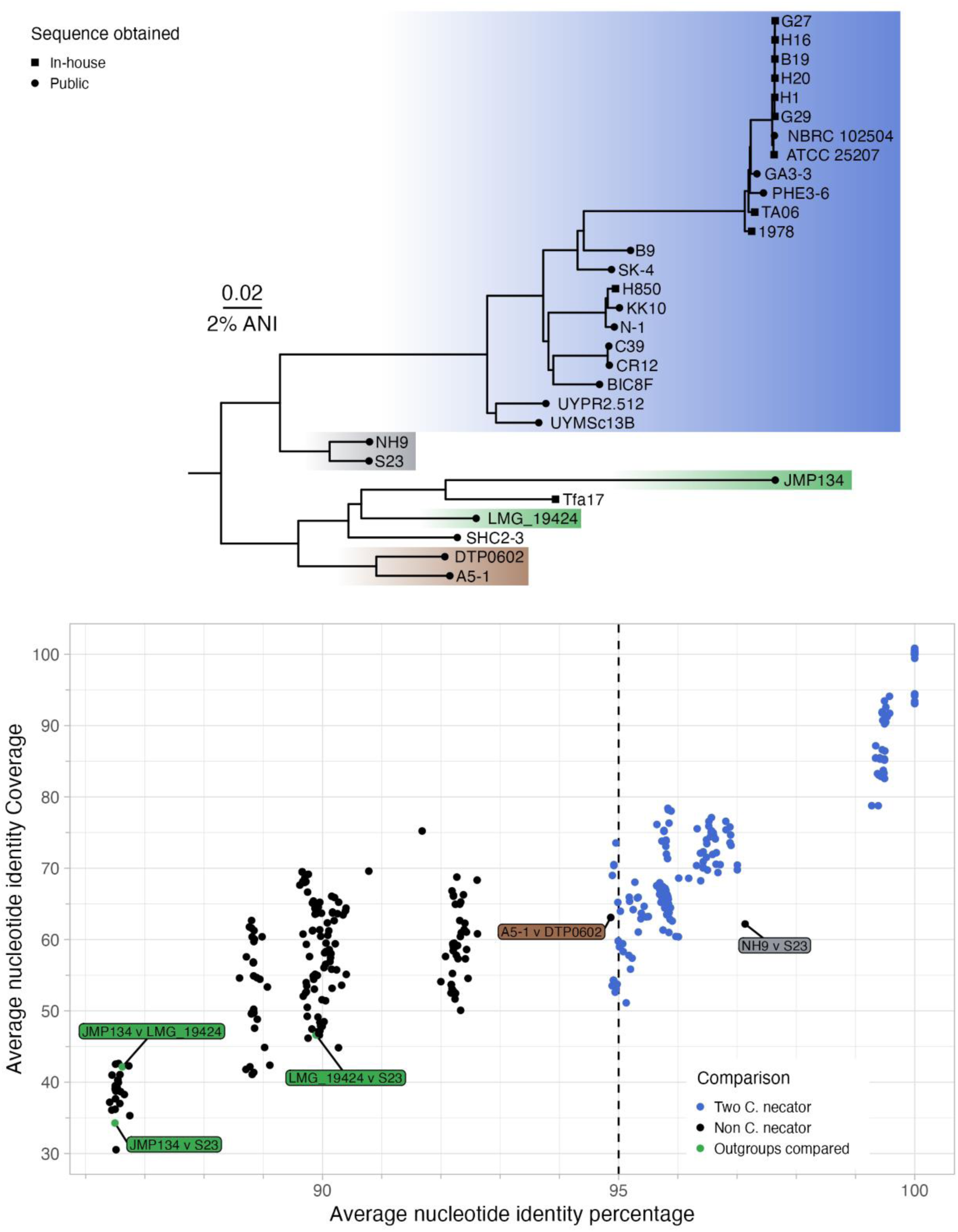
Average Nucleotide Identity (ANI) of refined *Cupriavidus necator* genomes and known outgroups, *cupriavidus lacunae (S23), cupriavidus pinatubunensis*i (JMP134), *cupriavidus taiwanensis (LMG 19424)*. A midpoint rooted neighbor joining tree constructed from ANI with tip labels indicating source of genome (public database or in-house sequencing) (**A**). The root edge was added for illustrative purposes. Colored rectangles mark interesting clades of the tree that are either one suspected species (Blue, *C. necator*. Grey, *C. lacunae*. Brown, *unknown*) or two known outgroups (Green). **B** illustrates the identity percentage (horizontal axis) and coverage percentage (vertical axis) of ANI comparisons of genomes. Colors indicate groups of compared strains, as given by the legend. Specific comparisons of interest, related to outgroups and other species, are highlighted by labels with strain names. A vertical dashed line indicates the common 95% ANI percentage cutoff for a species.

### High similarity between historical isolates

During the ANI analysis multiple genomes were identified as highly similar to strain H16. These genomes are mainly historical isolates with likely complex histories of documented and undocumented isolation, sharing between laboratories, and poor general traceability of handling. In an effort to document differences between strains we constructed a pangenome graph using pggb (36) of complete genomes with high ANI (>99%) compared to H16. Polymorphisms relative to H16 were identified (Supplementary Figure 3), revealing a low number of single nucleotide polymorphisms (n=[8,15]), deletions (n=[1,6]), insertions (n=[2,4]), and multiple nucleotide polymorphisms (n=[0,3]) across all isolates (Supplementary Table 2). As many of the historical strains were found similar to H16, we choose to exclude H1, H20, G27, G29, and B19 from further analyses. ATCC 25027 was included due to it not carrying the megaplasmid, pHG1, and NBRC 102504 was included due to it being a partially assembled genome.

### Genes of interest to metabolic functions of *Cupriavidus necator*

*C. necator* H16 is a strain well known for carbon dioxide fixation using the Calvin-Benson-Bassham (CBB) cycle, formate assimilation, hydrogen assimilation, nitrogen respiration, and PHA production. In addition, N-1 has been found to encode a methanol dehydrogenase (37). We searched for genes associated with these hallmark traits of *C. necator* to examine their presence across the species (Figure 2). Across all genomes we identified the presence of genes encoding functions related to the CBB cycle, utilization of formate, NO and NO_3_^-^ respiration, and PHA synthesis. These traits seem partially present in genome UYMSc13B, however the assembly of this genome is highly fragmented hindering identification of genes. *nisSCFDGHJNE* genes associated with NO_2_^-^ respiration was missing in the two monophyletic isolates of C39 and CR12. In these isolates the *nis* genes are missing, with the region, encoding *nis* genes in H16, otherwise occupied by five genes unrelated to the *nis* genes (Supplementary Figure 4). The methanol dehydrogenase encoded by N-1 can be found in related strains, H850 and KK10, along with the monophyletic clade of C39 and CR12. The acquisition of the methanol dehydrogenase may be different between the groups, as the gene is located on the chromid of N-1, H850, and KK10, contrasting the plasmid encoded methanol dehydrogenase of the complete assembly of C39 (Figure 2).

**Figure 2.**
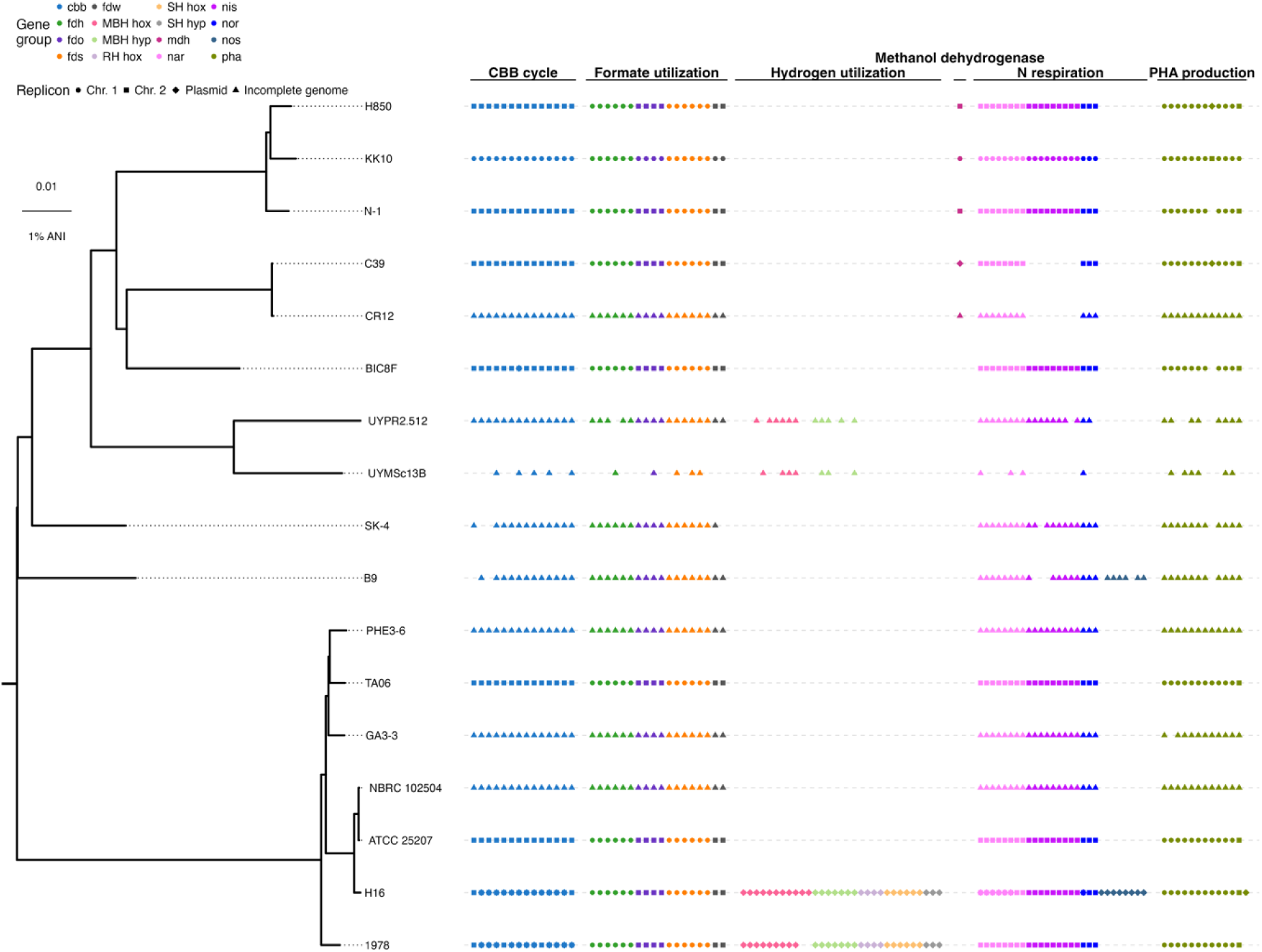
Presence and absence of genes related to hallmark metabolic functions of *Cupriavidus necator*. Relationship of strains is given by the midpoint rooted neighboring joining tree constructed from ANI percentage. Gene presence was determined using amino acids sequences of H16 or N-1encoded proteins, and identity and coverage cutoffs (see Methods). Points represent presence of genes with shape and color indicating related gene group and replicon placement, respectively.

Most sequenced strains do not contain genes for hydrogen utilization. An almost full set of homologs to the hydrogenases of H16 can be found only in strain 1978, with the exception of two membrane bound hox genes (*pHG004* and *pHG009*, *hoxR*)(Figure 3). The Soluble Hydrogenase (SH) and Actinobacterium Hydrogenase (AH) were found in 1978 with high similarities to the H16 versions. This is in contrast the Membrane Bound Hydrogenase (MBH) and Regulatory Hydrogenase (RH), which have lower similarities. As for H16 the hydrogenase genes in 1978 are encoded on a plasmid. The plasmid encoding hydrogenase genes in 1978 is seemingly not related to the pHG1 plasmid, of H16 (Supplementary Figure 5). Deeper analysis of the membrane bound hydrogenase of UYRP2.512 found *pHG004* and *pHG009* missing, similar to 1978. In addition, *hoxT* was missing and additional genes were inserted in the UYRP2.512 MBH region (Supplementary Figure 6). *hoxMLOQV* had low similarity to 1978 and H16, explaining their missing identification in the initial search of MBH genes (Figure 2). The *hoxJ* allele of H16 carries a mutation, G422S, resulting in a blocked autophosphorylation activity and a loss of regulation related to the presence of hydrogen (38). Aligning the HoxJ proteins of H16 and 1978 highlights a glycine in the conserved GxGL**G**L sequence of HoxJ. This indicates that 1978 should possess a fully functional RH, able to phosphorylate HoxA and regulate transcription of other hydrogenase genes.

**Figure 3.**
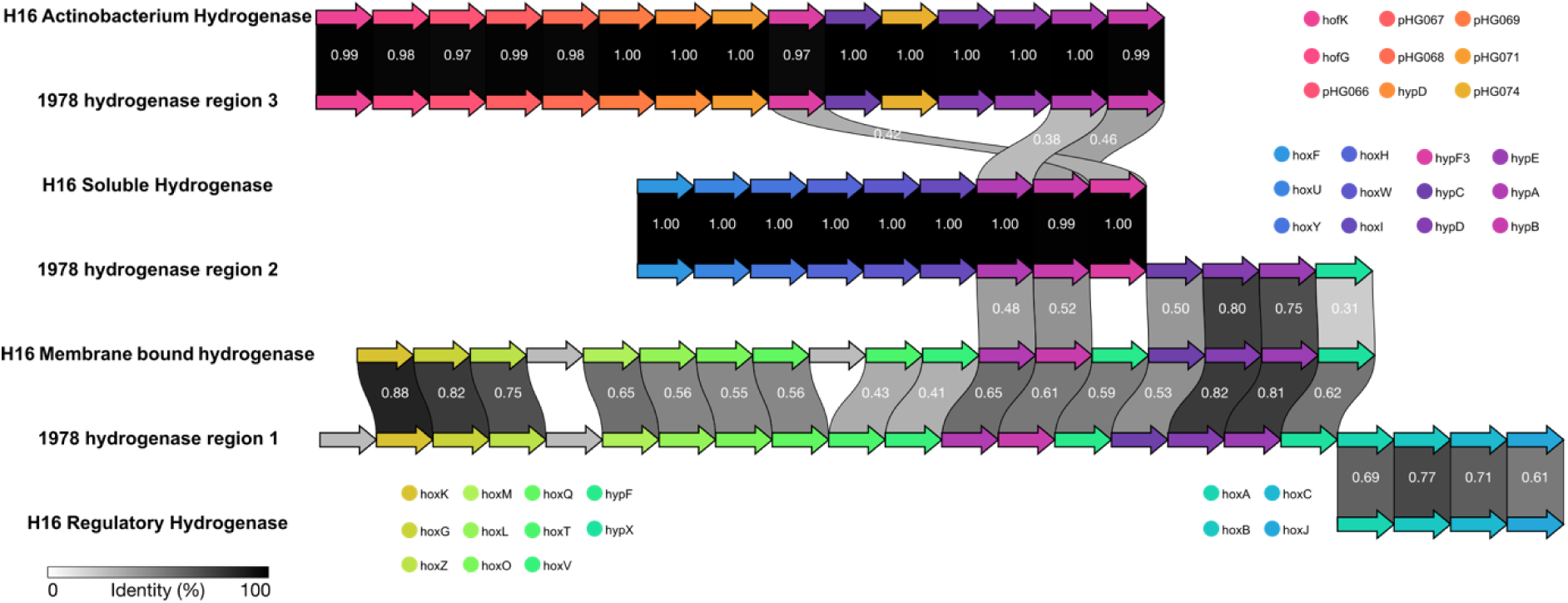
Clinker alignment of Hydrogenase related genes across strain H16 and 1978. Gene (arrow) size is not related to number of nucleotides. Color of genes is related to gene names, as given by the legend, with grey genes not being annotated or shared across strains. Links between genes indicate identity >30%, with specific identity indicated by grey scale color and number on links.

### Restriction modification systems and methylation motifs

Restriction modification systems can be a challenge when transforming bacteria, due to degradation of transformed DNA. To provide an overview of Restriction Modification (RM) systems in *C. necator* strains, we searched for these using Padloc, DefenseFinder, and hmm motifs via rmsFinder (39–42). The most common type of RM system was Type I, with 9 different systems identified across 13 strains (13/17, 76%)(Figure 4A). The second most common system was Type IV (n=7, across 5 strains)(Figure 4A). Both of the identified Type I and Type IV systems in H16 have previously been identified, acting as a positive control (9). The two RM systems of H16 are variably encoded genes, not being found as a core part of *C. necator* genome. As expected, the H16 RM systems are present in the closely related strains ATCC 25207 and NRBC 102504 (Figure 4A). As removal of RM Type I systems has been found to increase transformation efficiency in strain H16 (9), it is likely that removal of the identified genes could be used as a strategy to improve electroporation efficiency in the other strains. Carriage of DNA methylation motifs recognized by RM systems has also been shown to explain changes in transformation efficiency of plasmids (8). We reanalyzed our in-house Nanopore sequencing to identify methylated motifs of selected strains of interest, using Dorado, Microbemod, and STREME (43, 44). Methylation motifs in H16 have previously been examined using PacBio sequencing, fully determining the Type I system motif and an additional motif not related to an RM system (7). Using Nanopore sequencing we were able to identify both of the previously determined motifs. Similarly, in the closely related strain ATCC 25207, carrying the same set of RM systems, we identified methylation motifs identical to the ones in H16. To act as a negative control, we analyzed TA06. This strain was found to not encode any RM systems, and equally did not display any methylated motifs, besides the motif GTWWAC associated with the orphan methyltransferase (H16_A2613), which is a core gene across *C. necator*, with this matching motif being omnipresent in strains examined (Supplementary table 3). Finally, we examined strain 1978 and H850 due to their differences from H16, and encoding of a seemingly similar Type I system, allowing for cross verification of motif identification (Figure 4A). The motifs identified in strains 1978 (ACCNNNNNNGATC) and H850 (ACCNNNNNNNTGAA) demonstrated some differences (Figure 4B). Upon closer examination of the three genes which encode the Type I systems of these strains, the genes encoding the specificity subunit displayed a marked difference in identity (Identity ∼53%), likely explaining differences in the recognized motifs (Supplementary Figure 7). A fourth protein was identified in the RM system Type I region. It encodes a DUF262 domain which has recently been linked with multiple functions related to foreign DNA defense systems (45).

**Figure 4.**
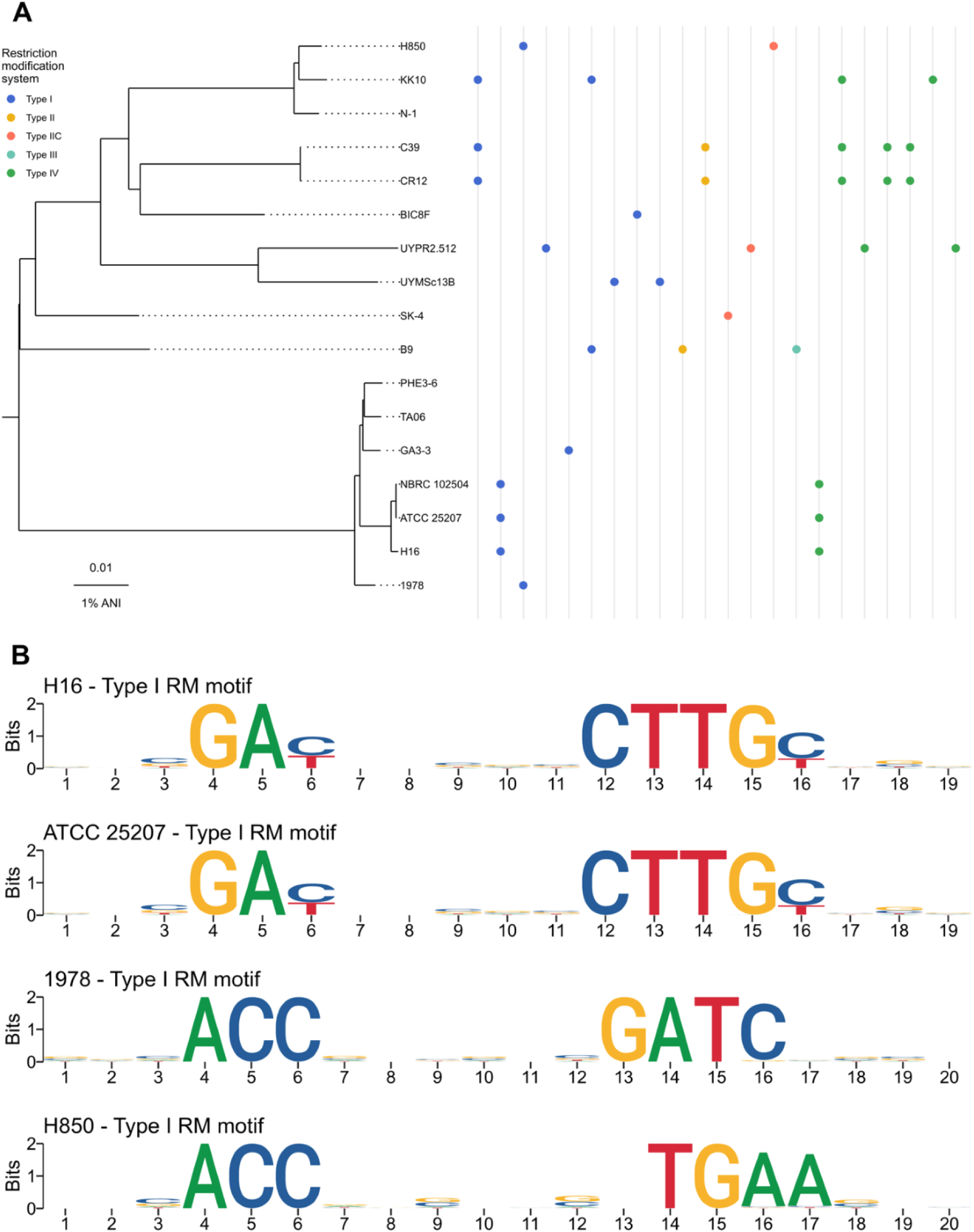
Restriction modification (RM) system of *Cupriavidus necator* strains. **A** illustrated presence of an RM system is indicated by a point with color specifying the type of system, as given by the legend. Each line indicate relation to a different set of RM proteins from databases used for search (see Methods). The phylogenetic tree is a neighbor joining tree similar to the one illustrated in Figure 2. **B** gives methylated motif identified for Type I RM systems across strains of interest.

## Discussion

*C. necator* is a species of increasing research interest, and an established model organism for aerobic hydrogen assimilation. The de-facto reference strain has over time become H16, but little work has been done to establish the genomic potential of other strains. We have provided a set of 22 genomes curated for species misannotations.

Having curated species designation of genomes, we have identified strains previously associated with *C. necator* that have been misannotated. Misannotation of species is not an unknown phenomenon but can impact genomic analyses and conclusions drawn in these. In addition to identifying misannotated genomes, we propose *C. lacunae* as a currently important outgroup for *C. necator*. *C. lacunae* is closely related to *C. necator*, consequently leading strain NH9 to be misannotated. With the identification of *C. lacunae* as a closely related species to *C.* necator, the ANI of 90% previously proposed for *C. necator* seems to be refutable. After performing comparisons among *C. necator* isolates, we observed a 94% ANI cutoff with a minimum of 50% coverage to be sufficient to capture genomes of interest. We do not propose these two cutoffs as strict cutoff but provide a collection of curated genomes along with appropriate outgroups that in the future could be used to better classify *C. necator* strain from other species.

The genomes of strains A5-1 and DTP0602 are indicated by ANI (94.9%) to be the same species. However, no known outgroup of this study or k-mer based taxonomic classification used were able to assign these strains to a species. The same is true for SHC2-3, no outgroup associated with this strain and no taxa was identified using a k-mer analysis. In contrast, strain Tfa17 from DSMZ seems more closely related to *C. oxalaticus* than to the annotated *C. necator,* based on k-mer based taxonomic classification. Verification and assignment of species for A5- 1, DTP0602, SHC2-3, and Tfa17 was outside the scope of this study, and thus remains an open questions.

In curating genomes deemed as *C. necator*, we identified low diversity among most historical isolates (H1, H16, H20, B19, G27, and G29) with ATCC 25207 and NBRC 102504 included. The identity of historical isolates observed here is unusual for strains that were supposedly isolated in separate events, some even in separate continents (H16 and H1 in Europe and H20 in North America)(46).

With the genomic insights obtained from newly sequenced genomes, we can propose three strains of increased interest, strain ATCC 25207, TA06, and 1978. These strains are similar to H16, but interesting based on current observations in the literature and for separate reasons. The genome sequence of strain ATCC25207 is separated from H16 by 11 single nucleotide polymorphisms, 6 deletions, and 2 insertions. The major difference between ATCC25207 and H16 is the absence of the megaplasmid, pHG1, in ATCC25207. This is similar to the ΔpHG1 mutant of H16 found to have improved growth on formate and multiple organic carbon-sources (47). Consequently, strain ATCC25207 may serve as a megaplasmid-free natural variant of H16. Along with ATCC25207, strain TA06 is the only other complete genome where no plasmid is identified. To this end these two strains can serve as megaplasmid-free chassis of *C. necator*.

Strain 1978 is interesting due to its regulation of hydrogenase gene transcription. H16 carries a HoxJ mutation (G422S) rendering its regulation of hydrogenase genes ineffective. This mutation is not present in 1978, indicating effective regulation of the hydrogenase related operons. To this end H16 is a natural overproducer of hydrogenase genes, taking up a sizable part of the proteome (48) and causing burden (47, 49). In Contrast, 1978 is hypothesized to be better at adapting to presence and absence of molecular hydrogen and unburdened by constitutive hydrogenase gene expression. Strain 1978 may therefore serve as an alternative to H16 when hydrogen assimilation is a desired trait, but the burden of constitutive hydrogenase expression is undesired.

All three strains, ATCC250207, TA06, and 1978, are available at DSMZ, and thus form a good base for future *C. necator* research. To complement their recommendation, we also provided methylation motifs for these strains to ease future genetic engineering campaigns.

Reassessing publicly available genomes of *C. necator* and presenting eight new and one improved genome assembly, we have been able to refine annotations of *C. necator* genomes and provide a curated collection of 22 genomes. Further, we have assessed hallmark metabolic traits for *C. necator* and can through this genomic analysis propose ATCC 25207, TA06, and 1978 as strains of interest for future research. Finally, we repurposed Nanopore sequencing data to identify restriction modification systems to aid future molecular biology campaigns examining above mentioned strains.

## Materials and Methods

### Data acquisition

Public genomes were identified in March of 2024 by using multiple strategies. Curation of NCBI Sequence Read Archive (SRA) and Assembly with search string: *("Cupriavidus necator" OR "Hydrogenomonas eutrophus" OR "Ralstonia eutropha" OR "Wautersia eutropha" OR "Alcaligenes eutrophus")* or *("Cupriavidus" OR "Hydrogenomonas" OR "Ralstonia" OR "Wautersia" OR "Alcaligenes")*. Assembled collections of publicly available reads with species annotations were interrogated (14). Genomes associated with *Cupriavidus necator* as a species on NCBI were collected on March 5^th^ 2024, removing duplicate genomes from a single strain. Finally, literature was searched for genomes previously associated with *C. necator* and appropriate outgroups.

### In House genome sequencing

Genomes ordered from DSMZ were grown in LB for DNA extraction using the GeneJET Genomic DNA Purification Kit (Thermo Scientific Inc.), following the manufacturer’s protocol. The quality of the extracted DNA was assessed using a 0.8% agarose gel, a Qubit 3 Fluorometer, and a NanoDrop 200/200c Spectrophotometer. DNA was split and subjected to either NanoPore sequencing using a R10.4.1 flow cell on a GridION, following a protocol optimized for high GC Actinobacteria genome sequencing (50), or sent for short read paired-end 350bp Illumina sequencing on a NoveSeq X Plus at NovoGene using a 6 cycle PCR protocol.

NanoPore and Illumina reads were subjected to a hybrid assembly approach using Hybracter (13) v.0.7.3. Hybracter is a long read first assembly pipeline, which yielded complete genome assemblies for all readsets. Hybracter was run with a minimum chromosome size of 2.5 Mb and default settings.

### Assessment of genomes

Kraken2(32) v.2.1.3 was used for k-mer based taxonomic classification with the Standard-8 Database from 2024/1/12 (ISO 8601) and default settings. ANI comparisons were made using pyani (33) v.0.2.12 with the ANIm and maxmatch method. For complete genomes replicons were compared using mge_cluster (51) v.1.1.0 to identify unitig presence and absence of replicons. Jaccard similarities of unitig presence absence was calculated using vegan (52) v.2.6-8 in R (53) v.4.3.2. H16-like genomes were compared using a pangenome graph constructed using pggb (36) v.0.5.4 with the deconstruct subcommand in vg (54) v.1.56.0 used to call polymorphisms against H16 and visualized with circlize (55) v.0.4.16 in R. Bakta (56) v.1.9.2 was used to annotate genomes with a protein file from H16 (GCA_000009285.2), gram neagative and the complete flags indicated. Comparisons of gene across genetic regions were conducted using Clinker (57) v.0.0.30. Gene of interest for metabolic functions were identified using Abricate(58) v.1.0.1 with proteins associated to functions from H16 (GCA_000009285.2) or N-1 (GCA_000219215.1) with cutoffs set default to 90% identity and 85% coverage. For the methanol dehydrogenase a 98% identity and 98% coverage were used. For the membrane bound hydrogenase genes 40% identity and 75% coverage was used, and for the regulatory hydrogenase 40% identity and 80% coverage was used. Phylogenetic trees and points were visualized using ggtree (59) and ggtreeExtra (60).

### Restriction modification systems and methylation motifs

Genes associated with restriction modification systems were identified using three methods: Defensefinder (45) v.1.3.0, padlock (41) v.2.0.0, and rmsfinder (39, 40) v.1.0.0. Identified restriction modification systems were curated and summaries across tools. Genes were related across genomes using a pangenome build using Panaroo (61) with clean-mode: sensitive, removal of invalid genes, a length difference percentage of 95%, refind-mode: strict, and a family identity percentage threshold of 60%.

Methylation motifs were identified by basecalling Nanopore reads with Dorado v.0.8.2 from Nanoporetech (https://github.com/nanoporetech/dorado). Reads were basecalled with the models: dna_r10.4.1_e8.2_400bps_sup@v5.0.0, dna_r10.4.1_e8.2_400bps_sup@v5.0.0_6mA@v1, and dna_r10.4.1_e8.2_400bps_sup@v5.0.0_4mC_5mC@v1 to identify 6mA, 4mC, and 5mC methylations. Reads were aligned to their respective genomes using minimap2 (62) v.2.28. Methylated genome positions and identification of motifs were identified using call_methylation from MicrobeMod (44) v.1.0.5,relying on STREME (43) v.5.5.5. Motifs were refined using the methylated_sites.tsv file given as output from MicrobeMod with sequencelogos illustrated using ggseqlogo (63) v.0.2. Results from Dorado and MicrobeMod can be found on Zenodo (see Data availability)

## Data availability

Code for reproduction of figures and underlying data is available at Zenodo under doi: 10.5281/zenodo.15075198. Newly sequenced genomes are available under bioproject: PRJNA1199890.

## Acknowledgement

The authors gratefully acknowledge the computational and data resources provided on the Sophia HPC Cluster at the Technical University of Denmark (64). The authors would also like to acknowledge Tue Sparholt Jørgensen and David Lokjær Faurdal on knowledge sharing on Nanopore sequencing. This work was supported by The Novo Nordisk Foundation (NNF20CC0035580 and NNF14OC0009473).

M.G.J. and S.D. conceived the study idea. E.F.V. and M.A.S. performed genome sequencing. M.G.J. and E.F.V. performed the genome assembly and bioinformatics analyses. M.G.J. and S.D. wrote the manuscript. S.D. and L.K.N. supervised the study. L.K.N. acquired funding.

---

**Supplementary Figure 1.**
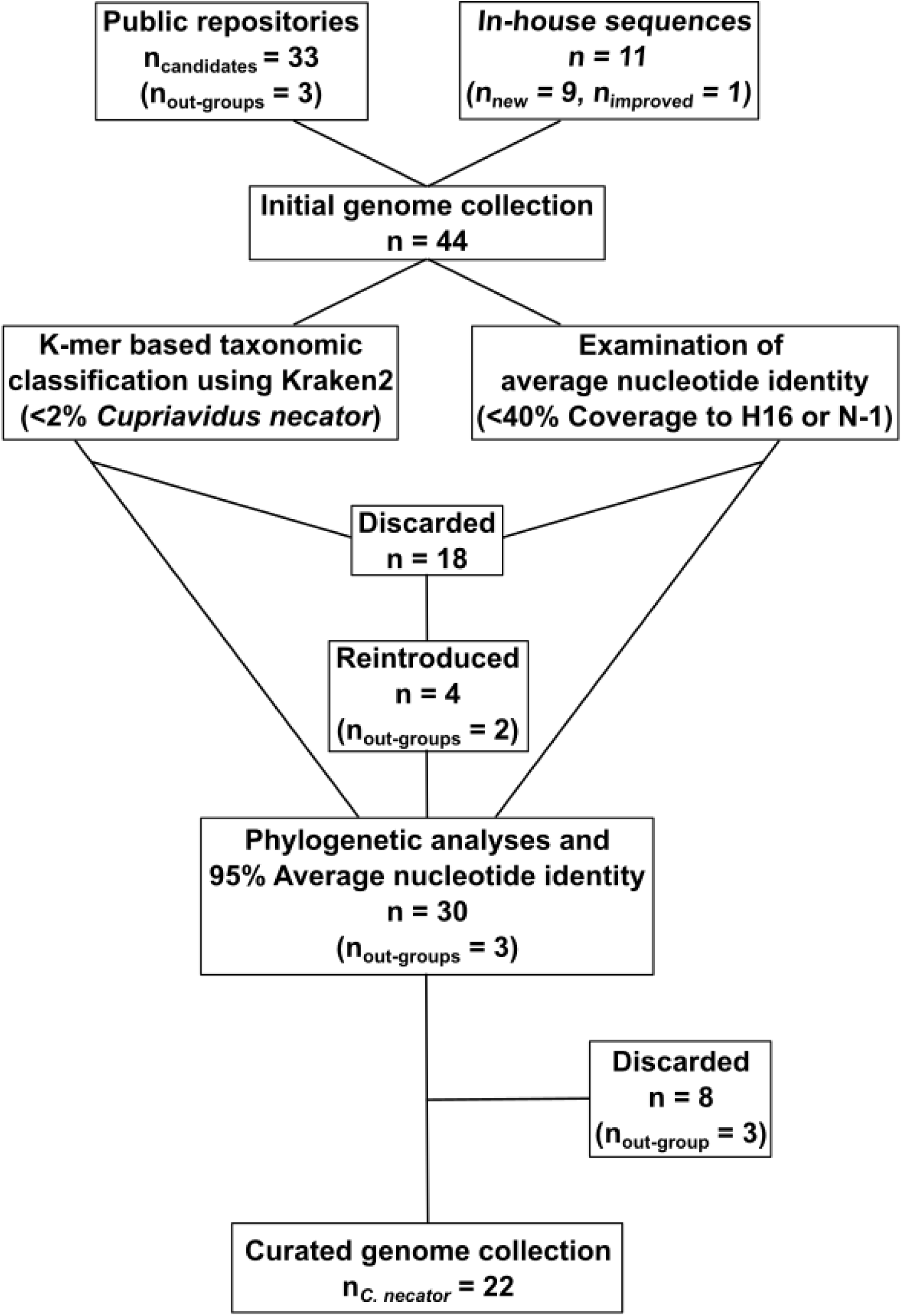
Flow of genomes from public repositories and in-house sequenced genomes to final curated genome collection. Each box is labeled with a step in the process of curation. For each step the number of genomes is given with a label “*n =* ”. “*n =* ” labels in parentheses indicate a subset of the above label and indicate numbers for specific groups of genomes that are of interest.

**Supplementary Figure 2.**
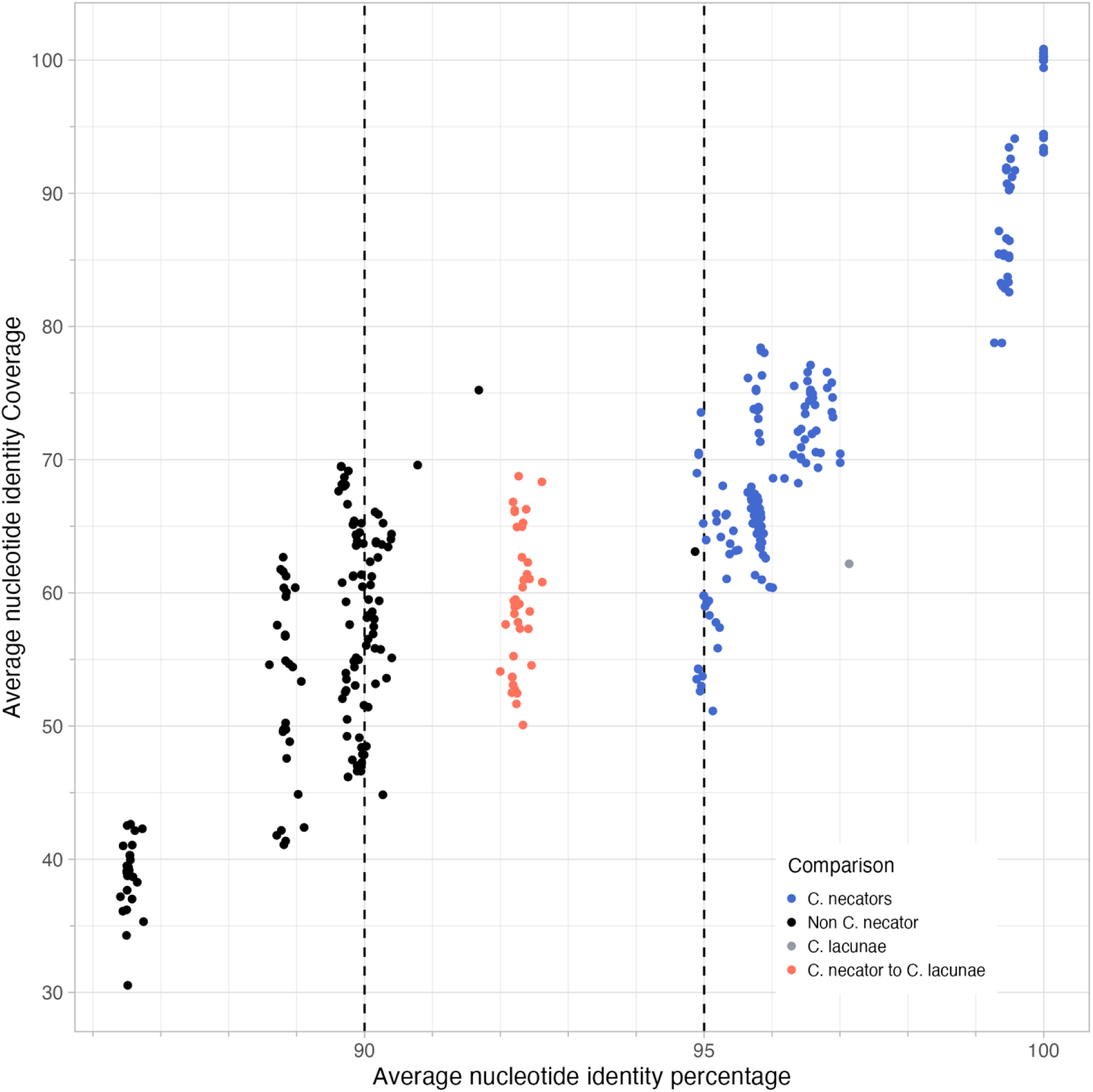
Illustration of Average Nucleotide Identity (ANI) percentage and coverage across pairwise comparisons of *C. necator*, *C. lacunae*, and non *C. necator* strains, colors indicated by legend. Vertical dashed lines indicate 90% and 95% ANI percentage.

**Supplementary Figure 3.**
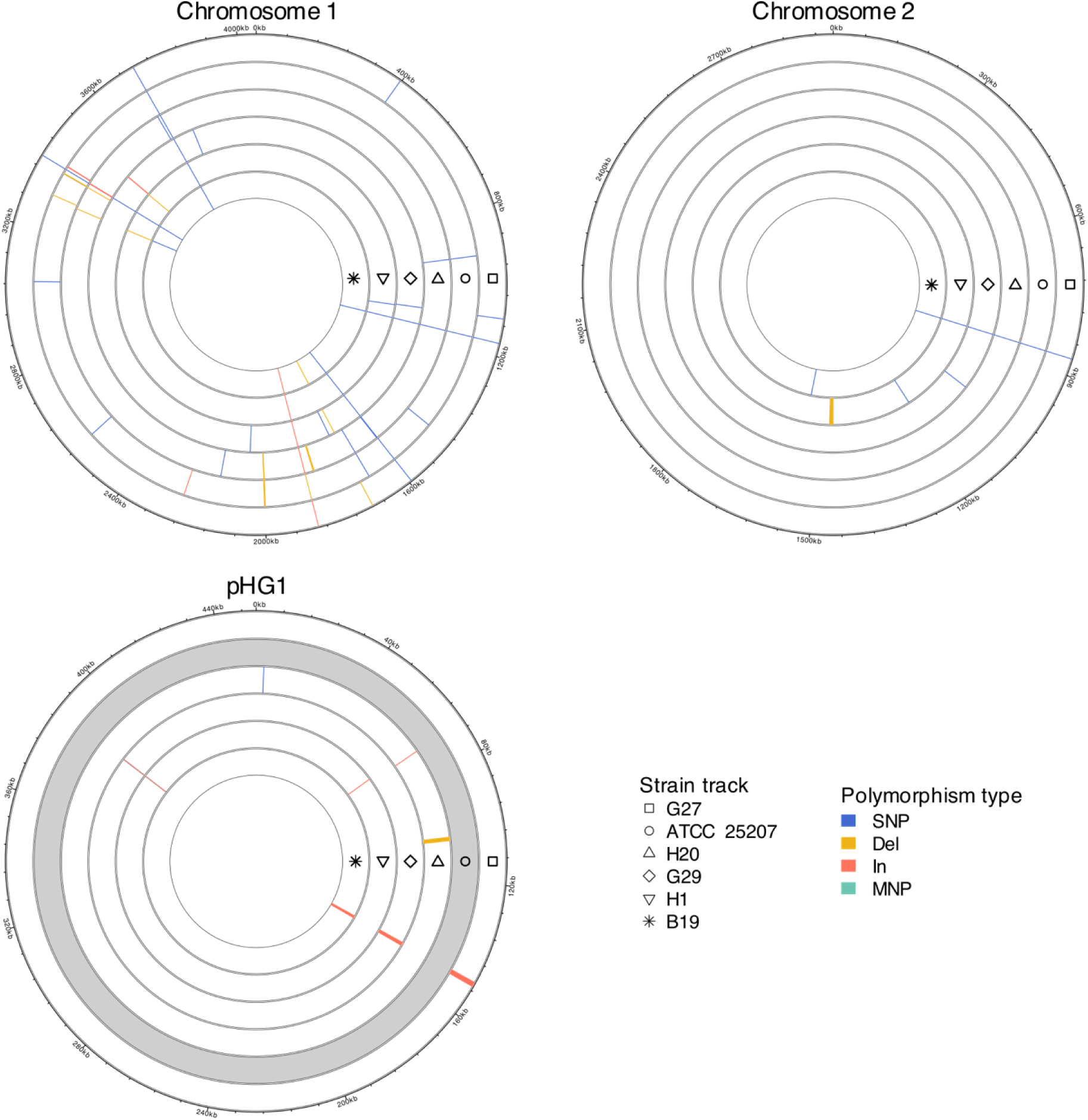
Comparison of replicons across strains found similar to H16. Each track of a circle represents a replicon from a genome, given by symbol at the 90-degree position and noted by the legend. Polymorphisms of a given, relative to strain H16, are marked by a color indicating the size and type of polymorphism, as given by the legend. Grey tracks indicate no presence of a given replicon.

**Supplementary figure 4.**
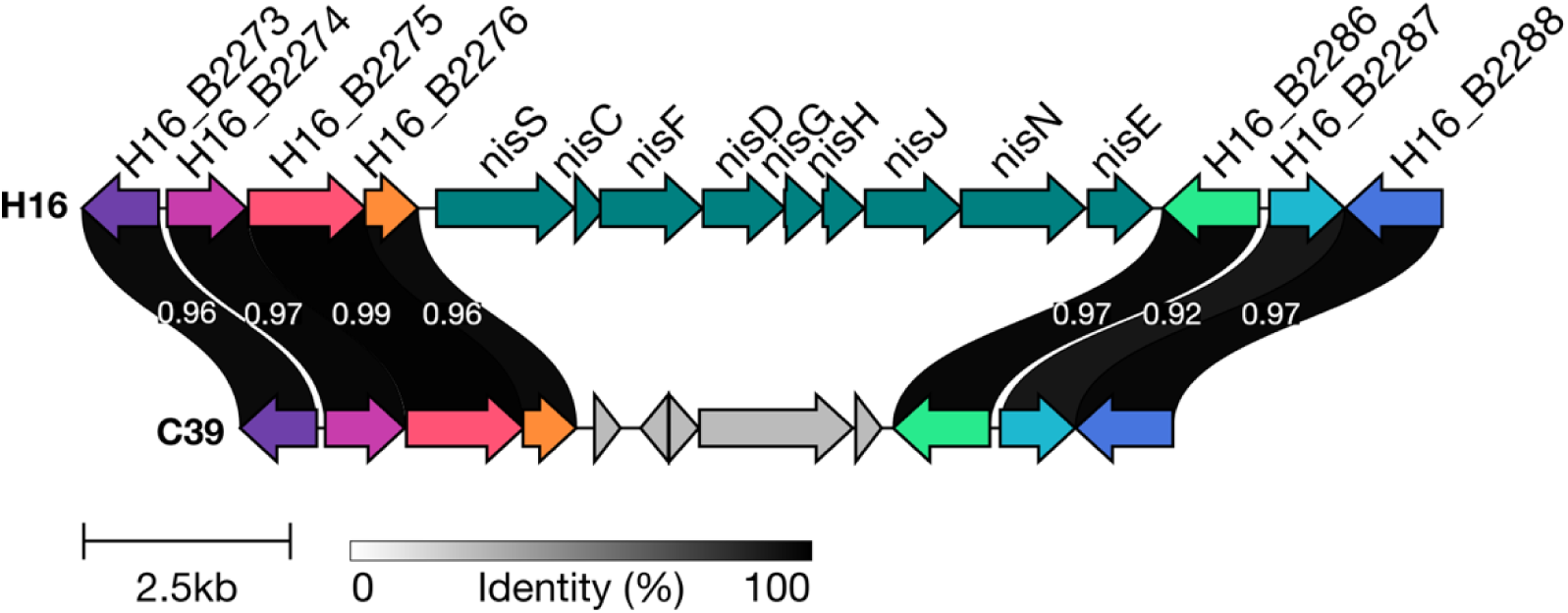
Clinker alignment of the *nisSCFDGHJNE* region of strain H16 and C39. Up- and down-stream genes are included and colored by match across strains. Links connect matched genes and indicate their identity by grey scale color. H16 Locus_tags of surrounding genes with *nis* genes being named.

**Supplementary figure 5.**
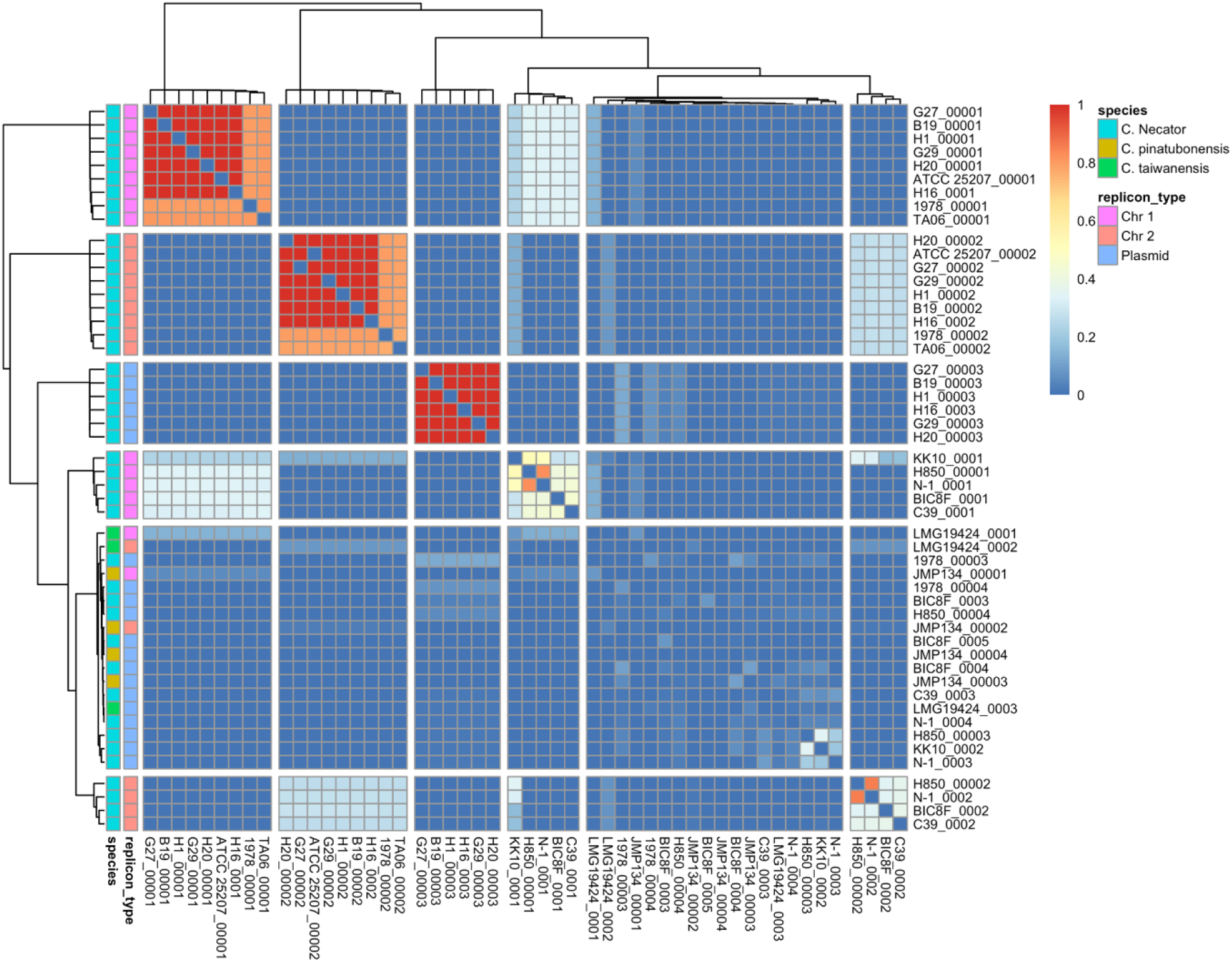
Comparison of replicons from complete *Cupriavidus necator* and two outgroup (JMP134 and LMG19424) strains based on Jaccard similarity of unitig presence and absence. Color of cells indicate jaccard similarity as given by red-blue scale. Species is indicated by left most column, followed by a column indicating the type of replicon (Chromosome, Chr, or Plasmid). Spaces among rows and columns were created by cutting row and column trees at n=6 groups.

**Supplementary figure 6.**
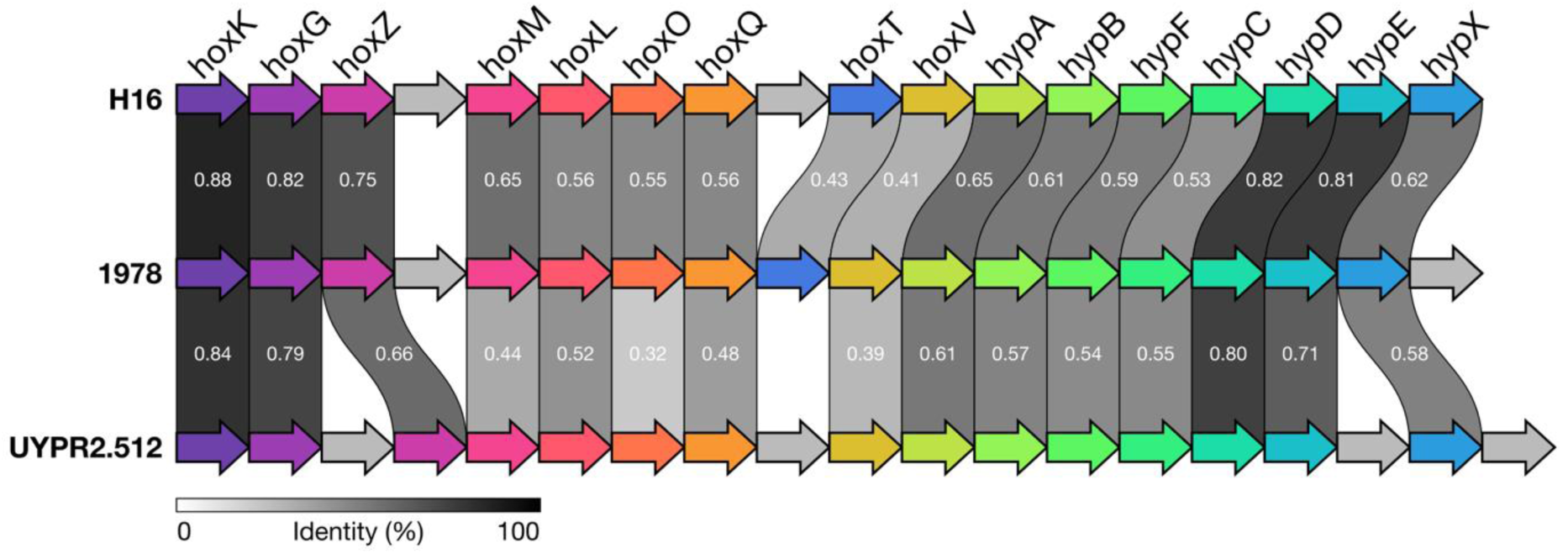
Clinker alignment of genetic regions encoding genes of the Membrane Bound Hydrogenase (MBH) of strains H16, 1978, and UYPR2.512. Arrows indicate genes, not to scale, with color indicating similarity across genomes. Grey arrows indicate no similarity for specific genes across genomes. Links between similar genes give identity (≥30%) of genes in grey scale. Name of genes are indicated above H16.

**Supplementary Figure 7.**
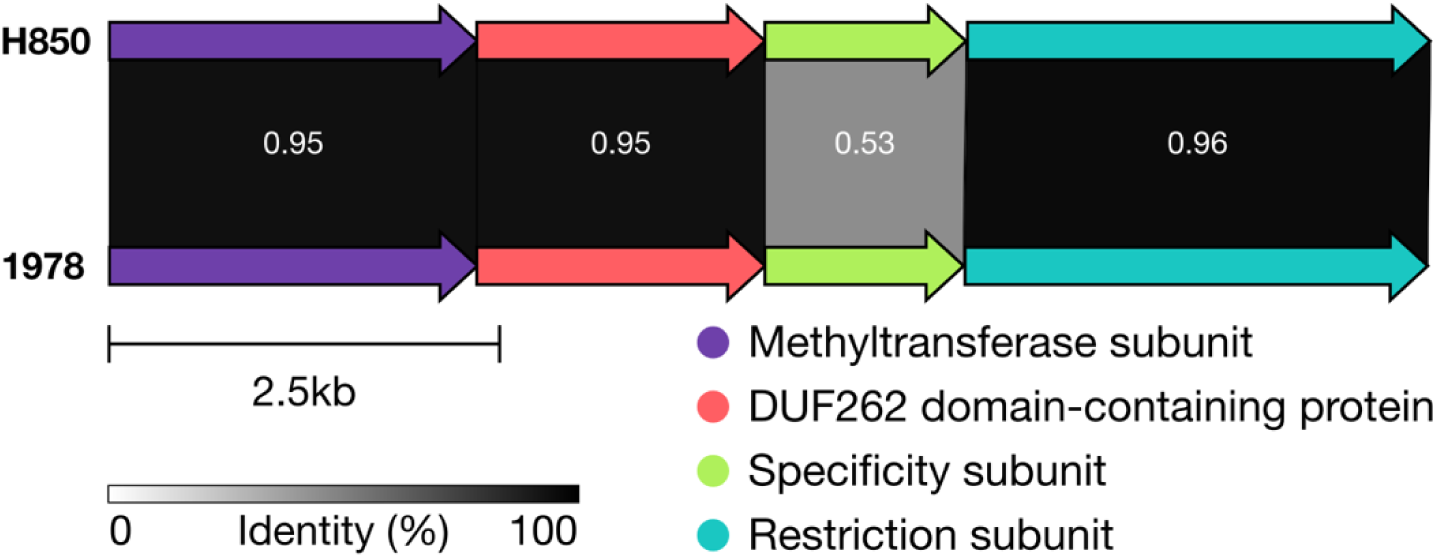
Clinker comparison of Type I restriction modification systems identified in strain H850 and 1978. Arrows indicate genes colored by similarity across strains and function. Links between genes gives the identity of protein sequences, as indicated by grey scale legend.

